# An RNA polymerase III tissue and tumor atlas uncovers context-specific activities linked to 3D epigenome regulatory mechanisms

**DOI:** 10.1101/2025.08.28.672650

**Authors:** Simon Lizarazo, Sihang Zhou, Ruiying Cheng, K C Rajendra, Yawei Shen, Qing Liu, Kevin Van Bortle

## Abstract

RNA polymerase III (Pol III) produces a plethora of small noncoding RNA species involved in diverse cellular processes, from transcription regulation and splicing to RNA stability, translation, and proteostasis. Though Pol III activity is broadly coupled with cellular demands for protein synthesis and growth, a more precise understanding of gene-level dynamics and context-specific expression patterns remains missing, in part due to challenges related to sequencing and mapping Pol III-derived small ncRNAs. Here, we establish a predictive multi-tissue map of human Pol III activity across 19 tissues and 23 primary cancer subtypes by comprehensively profiling the chromatin accessibility of canonical Pol III-transcribed gene classes. Our framework relies on the unique relationship between gene accessibility and Pol III transcription, inferring activity through uniform binary classification of ATAC-seq enrichment at Pol III-transcribed genes. By characterizing multi-context gene uniformity, we provide a definition of the core Pol III transcriptome, broadly active across specialized tissues, and catalog genes with varied levels of context specificity. Our genomic Pol III atlas uncovers variable levels of activity across tissues, including sharp contraction of the Pol III transcriptome in heart and brain tissues and frequent expansion across diverse cancers. We show that both tissue- and tumor-specific genes are significantly enriched within lamina-associated domains (LADs), and that aberrant expression of nuclear lamin proteins is sufficient to induce Pol III-emergent patterns at tumor-specific genes. Together, these findings link Pol III dynamics to subnuclear compartmentalization and provide a resource for better understanding Pol III expansion and small RNA biogenesis in cancer.

## Introduction

RNA polymerase III (Pol III) transcription is generally described as an essential housekeeping activity, producing small ncRNAs with central roles in diverse cellular processes, including transcription, RNA processing, and translation^1,2^. Importantly, Pol III transcription is found to be highly dynamic, and, for certain genes, its activity can be entirely restricted to one or a limited set of tissues^2–8^. Beyond examples of context-specific transcription, Pol III over-activity – a recurrent pattern during tumorigenesis – leads to the emergence of specific ncRNAs that are otherwise transcriptionally silent in most tissues^9–14^. These nuances contradict the simple housekeeping connotations often ascribed to Pol III, inspiring a deeper understanding of complex Pol III transcription patterns.

The multi-dimensional nature of Pol III transcription has remained difficult to measure, however, due to unique challenges and roadblocks to accurately defining Pol III activity. For one, the complex post-transcriptional RNA processing and modification events of small, highly structured ncRNAs require careful technical and bioinformatic consideration during library preparation and analysis stages, respectively^15–17^. Selective isolation of nascent Pol III-derived RNAs requires special steps, for example, including enriching for small RNA size distributions and accounting for the uncapped, 5’-triphosphate signatures of nascent Pol III products. The downstream analysis is further complicated by the multi-copy nature of many Pol III-transcribed genes, obstructing confident alignment of non-uniquely mapping reads^18^. While such challenges may not preclude a broad analysis of small RNA dynamics, they nevertheless present a challenge for defining gene-level transcription dynamics. These hurdles also undermine the utility of basic small RNA-seq experiments that are currently available through multi-tissues resources, such as ENCODE, GTEx, and others, for defining Pol III dynamics^19,20^.

Here, we circumvent these roadblocks by exploiting chromatin accessibility at Pol III target genes as a proxy for Pol III activity or inactivity, thereby overcoming the myriad challenges with profiling and mapping small RNA. We show that ATAC-seq (assay for transposase-accessible chromatin^21,22^) measurements scale with Pol III recruitment signatures and recover previously established tissue-specific patterns. In addition to improved read mappability, ATAC experiments cut directly to gene-level activity signatures and thus obviate challenges of interpreting transcription at the steady-state RNA levels. Overall, our genomic catalog of shared and context-specific Pol III transcription reveals a remarkable dynamic range of restriction, expansion, and an unexpected breadth of tissue- and tumor-specific Pol III activities. We further link these patterns with specific epigenomic regulatory mechanisms, including nuclear positioning of tissue- and tumor-specific genes within lamina-associated domains (LADs), altogether providing new insight on Pol III regulation and function in multiple contexts that are highly relevant to human health.

## Results

### A chromatin-based survey recovers context-specific Pol III transcription across human tissues

The Pol III transcriptome encompasses thousands of genes that encode various classes of small ncRNAs (Figure 1a), many of which themselves are comprised of variants (e.g. tRNA isodecoders) or repetitive gene copies (e.g. snaR-A ncRNA)^2,9,23,24^. We explored the relationship between chromatin accessibility for these various gene classes across human tissues with Pol III binding levels determined by available ChIP-seq data. Whereas genes absent Pol III are characterized by low accessibility, we find that ATAC-seq rises at sites with Pol III and further scales with increasing levels of Pol III ChIP-seq signal (Figure 1b). This broad relationship between Pol III occupancy and gene accessibility is consistent with our previous studies of Pol III dynamics in human monocytes, which established strong correlations between ATAC-measured chromatin accessibility with that of Pol III ChIP and nascent small RNA-seq levels within individual cells^25,26^. These observations suggest that gene accessibility alone may provide sufficient information to confidently predict whether a Pol III-transcribed gene is active or inactive in a particular context. We therefore developed a genomic framework to uniformly score ATAC-seq signatures across the Pol III transcriptome in human tissues. In brief, ATAC data corresponding to 19 distinct human tissues were optimally scaled to 250 million reads and filtered by saturation analysis (Supplemental Figure 1; see methods), and the observed signal intensities within annotated Pol III-transcribed gene classes were compared to a maximum, gene-centric local chromatin background (expectation, lambda) at 1, 10, 100 kb distances and whole-genome levels (Figure 1c)^27–29^. Genes of interest were thereafter defined as being active (“on”) or silent (“off”) depending on each gene-specific statistic.

**Figure 1.**
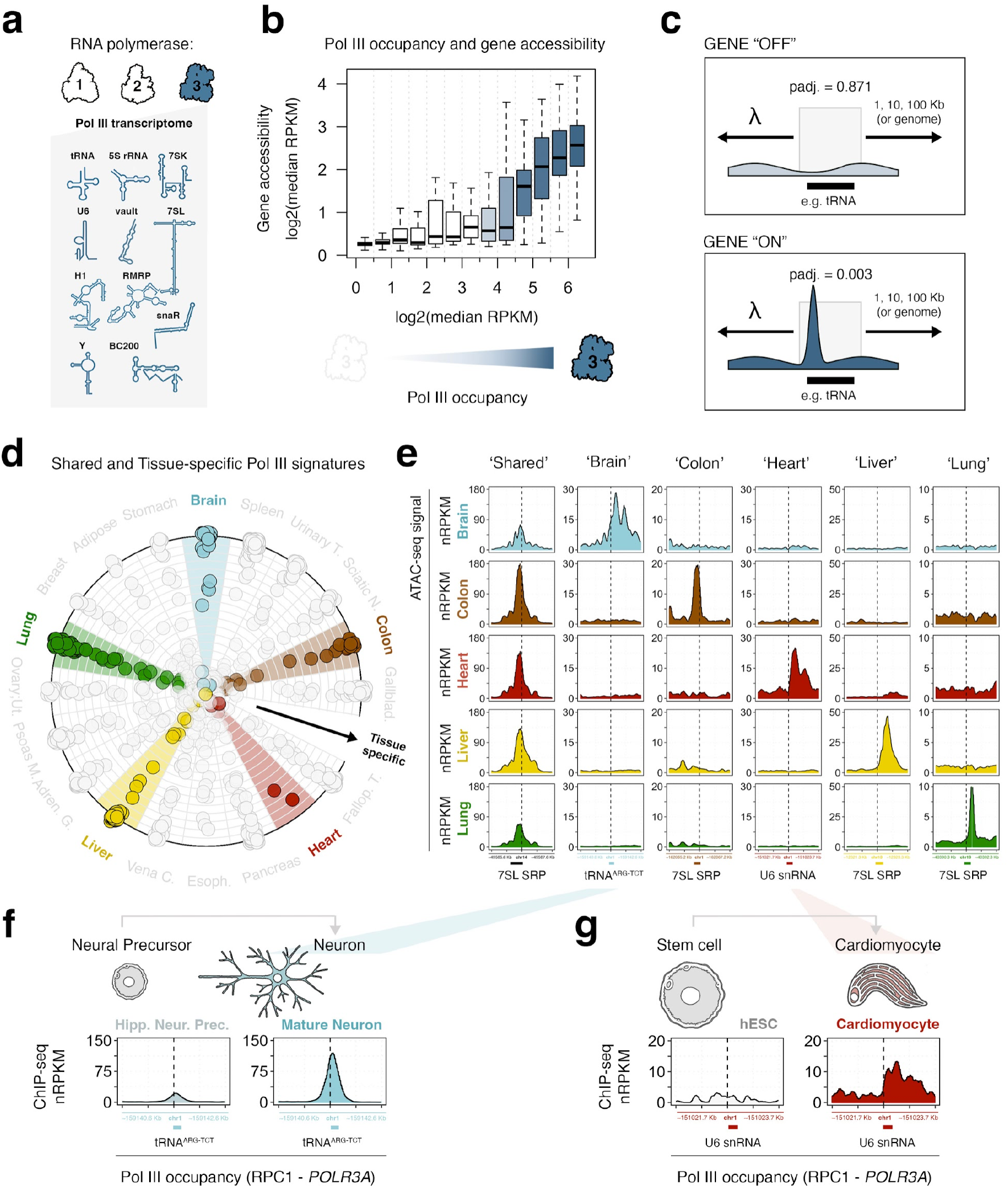
ATAC-measured gene accessibility scales with Pol III transcription signatures and recovers tissue-specific Pol III activity patterns. **(a)** Categorical overview of the small RNA species encoded by Pol III-transcribed genes and included in our large-scale genomic survey. **(b)** Analysis of chromatin accessibility (ATAC-seq) across Pol III-transcribed genes (shown in a), when binned by Pol III occupancy (ChIP-seq). **(c)** Illustrative overview of the gene “on” / gene “off” classification method used in this study. ATAC-measured gene accessibility is compared against a gene-specific local chromatin background (lambda); genes are inferred to be permissive and transcriptionally active (“on”) if accessibility signals are significant. **(d)** Visualization of tissue-specificity scores, measured by Shannon’s entropy (see methods). Individual data points represents Pol III-transcribed genes; radially positioned genes are characterized as context-specific. **(e)** ATAC-seq signal pileup at specific examples of Pol III-transcribed genes defined as shared (‘housekeeping’) or tissue-specific (‘brain’, ‘colon’, ‘heart’, ‘liver’, ‘lung’). nRPKM = normalized read per kilobase per million mapped reads. **(f-g)** Pol III occupancy at brain-specific and heart-specific genes (related to panel e). Pol III ChIP-seq data represent large subunit POLR3A (RPC1) binding in induced pluripotent stem cells (hiPSCs), hippocampal neural precursors, and mature neurons (f); or in human embryonic stem cells (hESCs) and cardiomyocytes (CM) (g).

To compare the resulting chromatin-based survey of Pol III activity across tissues, we further calculated an entropy-based uniformity score for individual genes, such that universally active genes are characterized by high categorical entropy, in contrast to tissue-specific genes marked by low entropy. Dominance-based visualization of multi-context gene entropy scores highlights examples of tissue-specific Pol III activity, where radially positioned genes reflect gene “on” status in a single tissue, and centrally positioned genes are shared by multiple or all tissues (Figure 1d). Inspection of several individual loci defined as having a shared or tissue-specific gene state demonstrates underlying ATAC patterns consistent with multi-context or context-restricted gene accessibilities (Figure 1e, Supplemental Figure 2).

We sought to validate the tissue-specific Pol III patterns predicted by our framework by investigating Pol III-binding in relevant cell types. First, we note that brain-specific Pol III signatures include accessibility of a tRNA-ARG-TCT encoding gene reported to be uniquely active in mature neurons^30^. ChIP-seq data for RPC1 (*POLR3A*) – the large subunit of Pol III – confirms robust occupancy in mature neurons (Figure 1f). Longitudinal characterization of Pol III localization in stem cell-derived cardiomyocytes CM^31,32^ similarly demonstrates examples of highly specific Pol III patterns. For example, a U6 snRNA gene marked by heart-specific chromatin accessibility is devoid of RPC1 signal in hESCs, with Pol III occupancy becoming evident specifically in differentiated CM (Figure 1g). These data help to further support the classification of heart-and other tissue-relevant Pol III signatures using gene accessibility, motivating a deeper analysis of the broader behaviors and dominant patterns of Pol III activity across human tissues.

### Tissue-level Pol III signatures point to variable levels of transcriptional restriction and expansion

Beyond simply capturing individual instances of shared or tissue-dominant signatures, we next explored whether gene status alone could collectively identify broader patterns of tissue-relevant Pol III activity. In support of this notion, hierarchical clustering of sample-specific gene signatures recovers tissue-level structure, including strong similarities among either brain, heart, liver, or lung-derived gene accessibilities (Figure 2a). These patterns, which are determined solely by the binary classification of gene state across Pol III-transcribed genes, indicate the presence of underlying patterns of tissue-specific Pol III transcription. This subclustering is further evident across nearly all the tissues surveyed in our study (Supplemental Figure 3). Overall, such patterns include a variable range of active Pol III-transcribed genes (i.e. total number) as well as specific subgroupings of genes that appear to be either activated or silenced within certain contexts. For example, tissue-level summaries of predicted Pol III gene states capture heterogeneous repertoires of active Pol III-transcribed genes, with smaller numbers in heart (195) and brain tissues (212) and comparatively larger gene sets in liver (308) and lung (357) tissues (Figure 2b-c). We find that tissues with expanded repertoires of active Pol III-transcribed genes naturally feature a higher number of context-specific signatures, whereas tissues with reduced Pol III repertoires are restricted to genes with a generally high level of uniformity across our multi-tissue survey (Figure 2b, Supplemental Figure 4).

**Figure 2.**
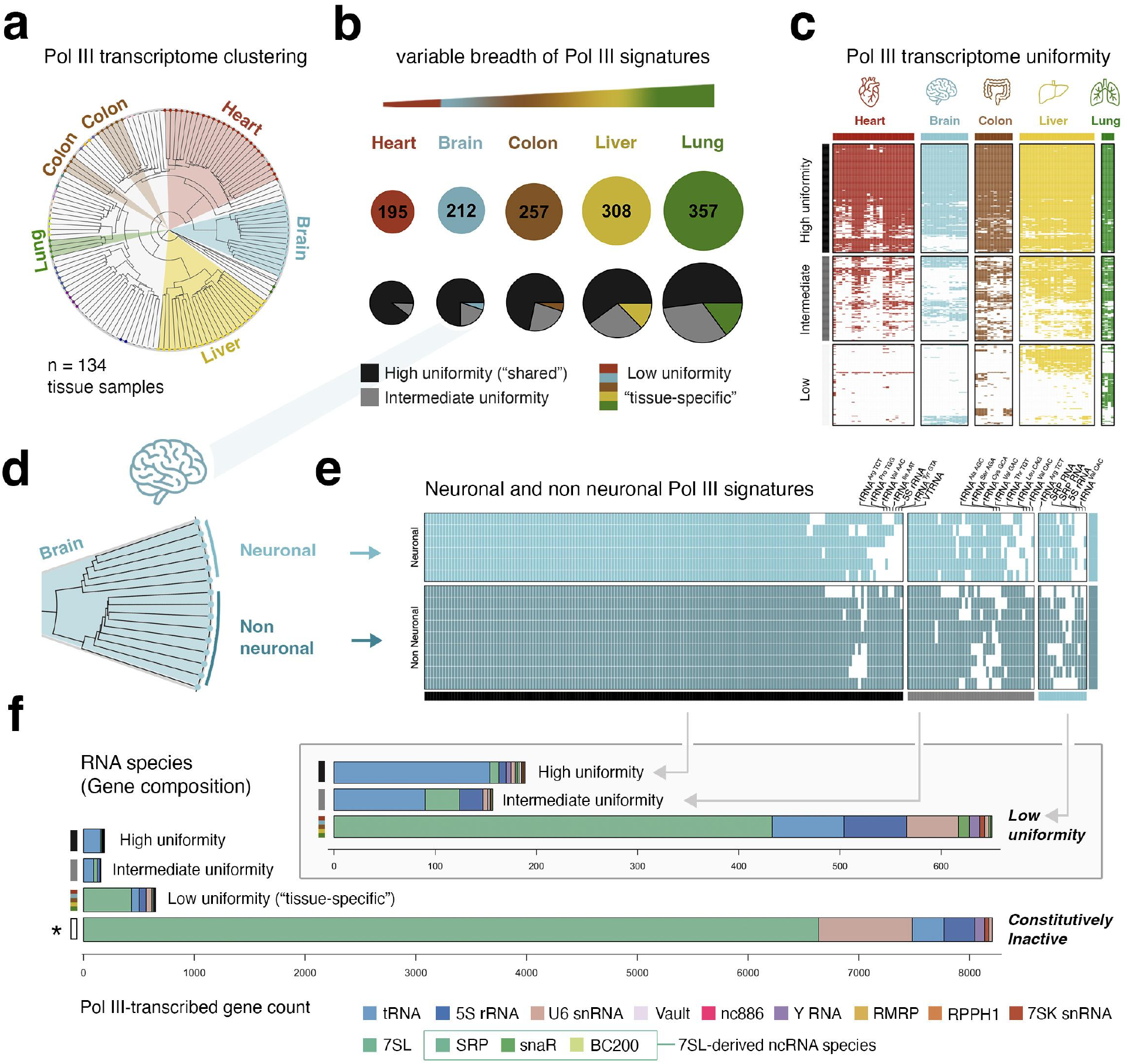
Tissue-level Pol III signatures include variable levels of restriction and expansion involving specific classes of Pol III-transcribed genes. **(a)** Hierarchical clustering of 134 sample-specific Pol III gene activity signatures. Individual experiments are categorically colored by human tissue, with the heart, brain, colon, liver, and lung highlighted for clarity. **(b)** Overview of the variable breadth and number of tissue-specific Pol III-transcribed gene features defined in the heart, brain, colon, liver, and lung tissues. **(c)** Visualization of Pol III-transcribed genes classified as having high, intermediate, or low uniformity and ensuing activity patterns within heart, brain, colon, liver, and lung tissues. Columns reflect individual experiments; tissue colors indicate “on” state, white color indicates “off” state. **(d)** Hierarchical subclustering of neuronal and nonneuronal samples derived from human brain tissues (related to panels a and d). **(e)** Heatmap visualization of 212 active Pol III-transcribed genes in human brain tissues in neuronal (light blue; top) and nonneuronal experiments (dark blue; bottom), subdivided by multi-tissue classification of high, intermediate, or low gene uniformity (related to panels b and c). **(f)** Composition of high, intermediate, and low uniformity subclassifications on the basis of annotated Pol III-transcribed gene subcategories. Inset serves to highlight these subcategories, in juxtaposition to the expansive number of annotated but transcriptionally inactive instances of various Pol III-transcribed gene classes (bottom barplot).

Even within tissues with comparatively reduced Pol III transcriptomes, our analysis further captures structures indicative of subtissue-level Pol III activities. For example, brain tissue data derived from neuronal and nonneuronal origin, which subcluster based on subtype (Figure 2d), reveal small shifts in predicted Pol III activity between these contexts (Figure 2e). These differences tend to involve neuronal-specific loss of activity at tRNA genes marked by relatively high and intermediate uniformity across tissues, as well as more subtle neuronal- and nonneuronal-specific activities at genes encoding tRNA, 5S rRNA, and SRP-related RNA species (Figure 2e). These patterns are consistent with the compositional bias found within each subcategory of tissue uniformity, such that genes with high and intermediate uniform activity generally encode tRNAs and other classes of essential RNA species, whereas genes with low uniformity are dominated by signal recognition particle (SRP)-related RNA species (Figure 2f). However, our survey also identifies well-defined Pol III-transcribed genes with highly variable activity signatures across the 19 tissues, such as *VTRNA1-2* and *VTRNA1-3* - vault RNA genes that are predicted to be active in multiple tissues (e.g. pancreas, gallbladder, colon, etc.) but silent in several others (e.g. brain, sciatic nerve, esophagus), exemplifying the information-rich nature of the multi-tissue Pol III atlas (Supplemental Figure 2).

### Sequence-based prediction of Pol III gene activity and tissue-specificity

Given the novel scope of our multi-tissue Pol III activity map, we next explored whether specific sequence elements could predict gene signatures, such as active vs. inactive or high vs. low tissue uniformity. However, given that Pol III-transcribed genes include heterogeneous promoter architectures and unbalanced frequencies, we restricted our analysis to tRNA genes, which near-universally depend on gene-internal A-box and B-box promoter elements^33–36^. To avoid introducing prior assumptions, we first performed *de novo* motif enrichment across the full set of active and inactive tRNA genes using STREME - a sensitive sequence analysis tool designed to capture overrepresented motif signatures^37^. Predictive values among the resulting set of enriched motifs, measured by mean decrease in accuracy (MDA) from a random forest classifier, were then ranked according to their ability to predict active tRNA genes from genes that are universally silent across all examined tissues (AUC = 0.7989, Supplemental Figure 5a). Notably, we find that the strongest predictor of tRNA gene state corresponds to a sequence with significant similarity to previously reported B-box elements (Figure 3a, Supplemental Figure 5b)^33,38,39^. In contrast, the sequence most powerful in distinguishing tRNA genes with intermediate and high uniformity (i.e. broad expression) from genes with low uniformity (i.e. tissue-restricted expression) aligns significantly to previously reported A-box elements (Figure 3b, Supplemental Figure 5b). These sequences appropriately map to the expected positional windows of A- and B-box elements within tRNA genes and, using the positional distribution of these features, we additionally recover empirical representations of the human A- and B-box elements derived from our multi-tissue atlas of active tRNA genes (Figure 3c-d).

**Figure 3.**
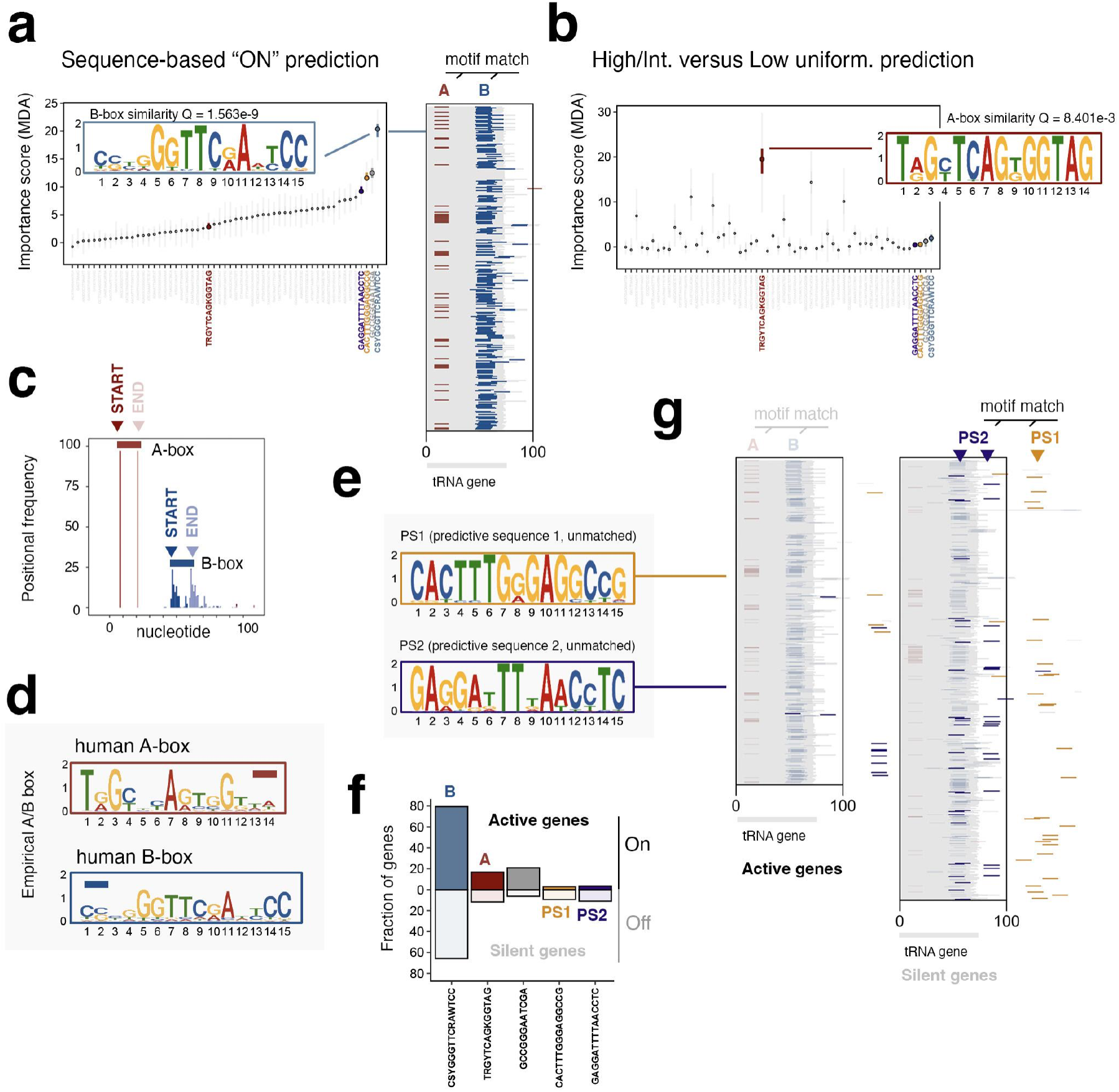
Sequence-based prediction of tRNA gene activity patterns recovers A- and B-box promoter elements, as well as uncharacterized gene proximal elements. **(a)** Predictive power of sequences enriched at tRNA genes for classifying active versus inactive tRNA genes. Motifs - discovered de novo by STREME - are ranked by Random Forest importance score. A 15-mer sequence element with maximum importance and significant overlap with previously reported B-box elements is highlighted. Sequences with significant B-box (blue) and A-box (red) similarity are further positionally mapped within active tRNA genes (inset). **(b)** Analogous plot of sequence elements ranked in panel a, but showing Random Forest importance score for predicting high or intermediate uniformity versus low uniformity (i.e. tissue-specific) tRNA genes. A 14-mer sequence element with maximum importance and significant similarity to previously reported A-box elements is highlighted). **(c)** Positional frequency histograms of A-box and B-box-like elements in active tRNA genes. **(d)** Empirical human A- and B-box sequence elements derived by positional frequencies shown in panel c. **(e)** Uncharacterized predictive sequences PS1 and PS2 shown to have high importance in classifying tRNA active versus inactive state. **(f)** Individual frequencies of predictive sequence elements within active and inactive tRNA genes. In contrast to other features, PS1 and PS2 occur at higher frequency in inactive genes. **(g)** Positional map of PS1 (gold) and PS2 (purple) occurrences within active and inactive tRNA genes.

In addition to the *de novo* B-box-related sequence, we discover multiple motif signatures with predictive power in distinguishing active and inactive tRNA gene patterns, including a slightly offset B-box sequence that likely extends from the B-box element itself (Figure 3a, gray). Two additional sequences have notable predictive power but are otherwise unannotated in terms of established regulatory sequences and transcription factor binding sites (Figure 3e). In contrast to both A and B box elements, these predictive sequences – coined PS1 and PS2 – are commonly observed in inactive rather than active tRNA genes, indicating a negative relationship with Pol III (Figure 3f). We also find that PS1 is positionally biased downstream of the annotated tRNA gene, whereas PS2 is located both within and downstream of certain tRNA genes (Figure 3g). It is possible that PS1, PS2, and/or variations in the A-box sequence itself modulates tRNA gene competence through interactions with transcription or chromatin factors. Though overlap enrichment analyses for available ChIP-seq data (ChIPatlas) fail to identify candidate factors linked specifically to PS2, we find that PS1 sequences – which occur in AluSC family SINE elements – have some level of enrichment corresponding to Lamin B1 and DDX5 ChIP-seq data (Supplemental Figure 5c-d)^40–44^. Overall, the presence of DNA-level indicators of tRNA gene activity supports a model in which gene-intrinsic features may contribute, in part, to tissue-specific Pol III transcription patterns^33^.

### Tissue-restricted Pol III-transcribed genes are enriched in nuclear lamina-associated domains (LADs)

Moving beyond gene-intrinsic sequence features, we next explored whether specific transcription factors, chromatin and DNA modifications, or 3-dimensional (3D) epigenome signatures were similarly predictive of Pol III gene activity. Our survey included currently available ATAC, ChIP, and DNA methylation data (ChIPatlas)^42–44^, as well as nuclear lamin DamID (4DNucleome)^45–47^ totaling ~30,000 experiments (Figure 4a). Multivariate analysis on 147 unique epigenomic variables is robust at discriminating Pol III-active genes (AUC = 0.86, Figure 4b), and identifies multiple features important for prediction, including H3K79me2, chromatin accessibility (as expected), and several TFs (Figure 4c). Nuclear lamin proteins - mapped by DamID - are notable among this list of chromatin factors, particularly because these patterns are stronger at Pol III-transcribed genes predicted to be inactive (Figure 4c-d).

**Figure 4.**
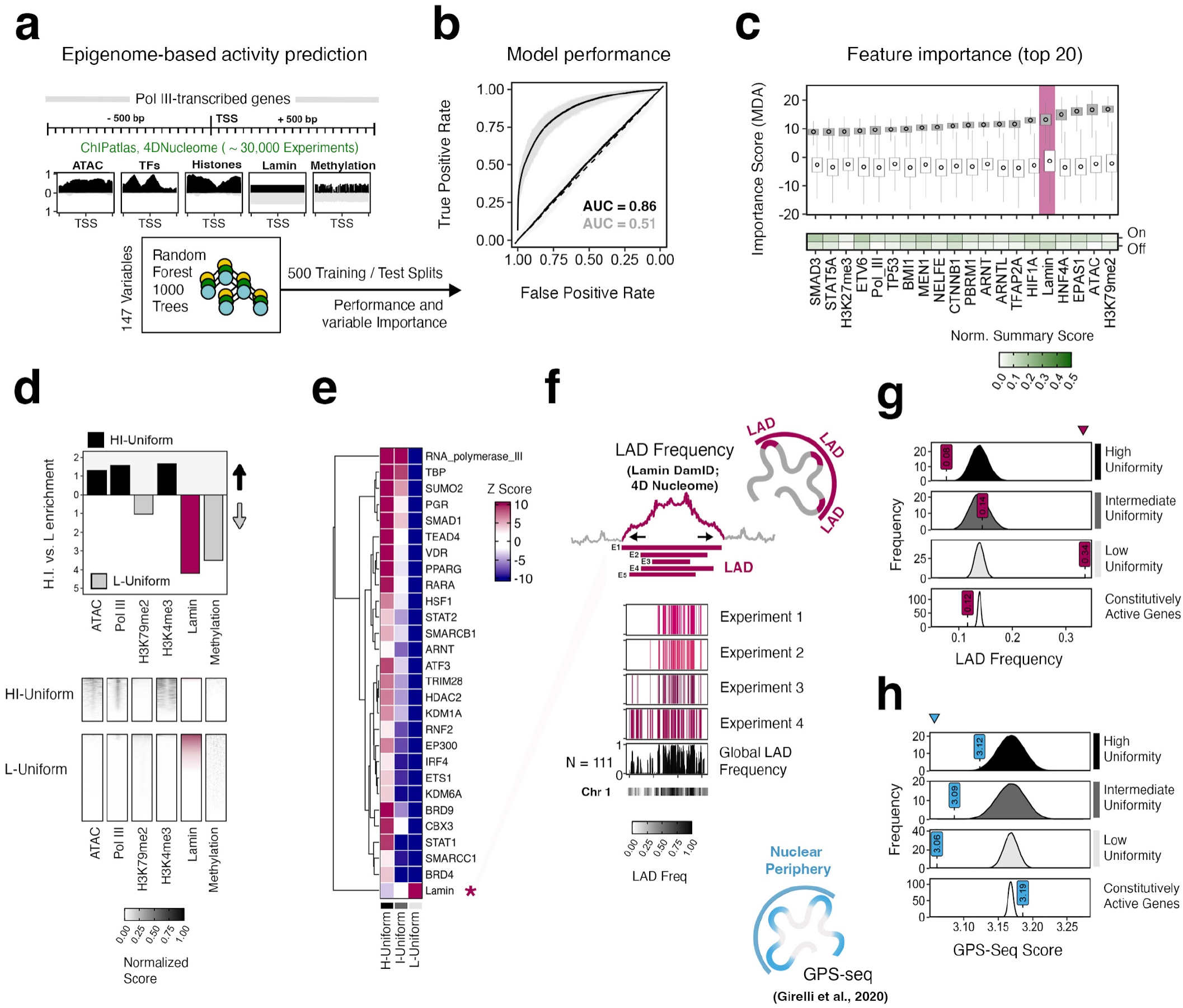
Epigenome-based prediction of Pol III activity uncovers strong enrichment of context-restricted ncRNA genes within Lamina-associated domains (LADs) and peripheral subcompartments. **(a)** Illustrative overview of the epigenome-based survey for features that predict Pol III gene activity. Random Forest analysis incorporated 147 features including chromatin accessibility (ATAC), Transcription Factor (TF)-binding, histone modifications (ChIP-seq), Lamin association (DamID) and DNA methylation (Bisulfite-seq) profiles retrieved from ChIPatlas and 4DN. In brief, intra-factor gene-level statistics spanning 1 kb gene-centered intervals were normalized across all Pol III-transcribed genes. Multivariate class prediction was then evaluated over 500 training-test splits (75 v. 25). **(b)** Classification model performance; Mean AUC (Area Under Curve) = 0.86; response variable randomized AUC = 0.51. **(c)** Ranked importance of epigenomic features for classifying Pol III-transcribed gene state across all human tissues (MDA = mean decrease accuracy). Only the top 20 features are shown for clarity; Normalized summary score represents average gene-level signal statistic across individual gene groups. **(d)** Comparison of feature signal enrichment at high and intermediate (H.I.) uniform genes versus low (L) uniformity genes. Barplot (top) depicts the relative enrichment for genes with higher-than-median signal. Heatmap (bottom) visualizes normalized signal scores at 10 bp resolution (bin size) for individual genes in each subgroup. **(e)** Heatmap of feature enrichment z-scores at high, intermediate, and low uniformity genes; only features with significant enrichment (z > 2 in one or more groups) are shown for clarity. **(f)** Illustrative overview of LAD frequency calculation, which takes into account all LAD (4DN Lamin DamID-based) calls and computes genomic cross-experiment frequency. **(g)** Observation and empirical null distributions for average LAD frequencies spanning high, intermediate, and low uniformity genes. **(h)** Analogous analysis corresponding to radial subnuclear positions mapped by GPS-seq.

To further reinforce our classification-based Pol III analysis, we additionally profiled the enrichment of all chromatin and transcription factors at Pol III-transcribed genes stratified by tissue-uniformity. Comparing the overall frequency of these factors at each gene group against a null distribution that considers active genes, our analysis appropriately captures Pol III-binding enrichment at both high and intermediate uniformity genes (Figure 4e). We notably detect significant enrichments for a multitude of other factors, including both established regulatory players (e.g. TFIIIB subunit TBP, Progesterone Receptor PGR^48^) and several potentially novel gene regulatory mechanisms. Conversely, only Lamin proteins are significantly enriched at low uniformity genes, reaffirming our prior classification results (Figure 4e, Supplemental Figure 6).

Lamin enrichment, which likely indicates peripheral enrichment at the nuclear lamina, mirrors that of DNA methylation, both of which are mechanisms generally associated with gene silencing (Figure 4d)^49–51^. We therefore further explored the differential enrichment of Pol III-transcribed genes within lamina-associated domains (LADs). In brief, a multi-context LAD frequency was first determined by integrating all currently available Lamin DamID-based LAD calls in humans (4DNucleome, n = 111 experiments), producing a genome-wide summary of LAD interactions (Figure 4f)^52,53^. Using a randomization-based framework, we find that Pol III-transcribed genes with high and intermediate uniformities are characterized by low LAD frequencies, as expected for active genes that are expressed in most or all tissue contexts (Figure 4g). In contrast, tissue-restricted genes with low uniformity scores are associated with significantly higher LAD frequencies, consistent with subcompartmentalization of tissue-specific genes at the nuclear lamina (Figure 4g). These findings are further supported by orthogonal measures of gene location relative to the nuclear periphery, queried by GPS-seq - a longitudinal restriction enzyme digestion method^54^. In these experiments, low uniformity genes are characterized by substantially lower GPS-seq scores, consistent with a gene position bias at the nuclear periphery (Figure 4h).

### Cancer-emergent Pol III signatures predominantly arise at tumor-specific, LAD-associated genes

We next investigated cancer-associated Pol III patterns by applying the same ATAC-based framework and entropy analysis to 328 quality-filtered experiments spanning 23 distinct primary tumor-types (The Cancer Genome Atlas)^55–57^. As described for tissues, we find that Pol III-transcribed genes in tumors are characterized by varied patterns, with certain genes actively shared across contexts and others restricted to a particular tumor type (Figure 5a). Of note, our cancer-associated Pol III survey also identifies more than 450 “new” genes that are predicted to be active in 1-or-more tumors but inactive in all normal tissues studied (Figure 5b-c). Upon inspection, we find examples of such cancer-emergent genes that are definitively tumor-specific (e.g., LUAD and LUSC examples; Figure 5b), whereas other instances are characterized by weaker, statistically insignificant accessibility in closely related tissues (e.g. colon and COAD; liver and LIHC examples; Figure 5b).

**Figure 5.**
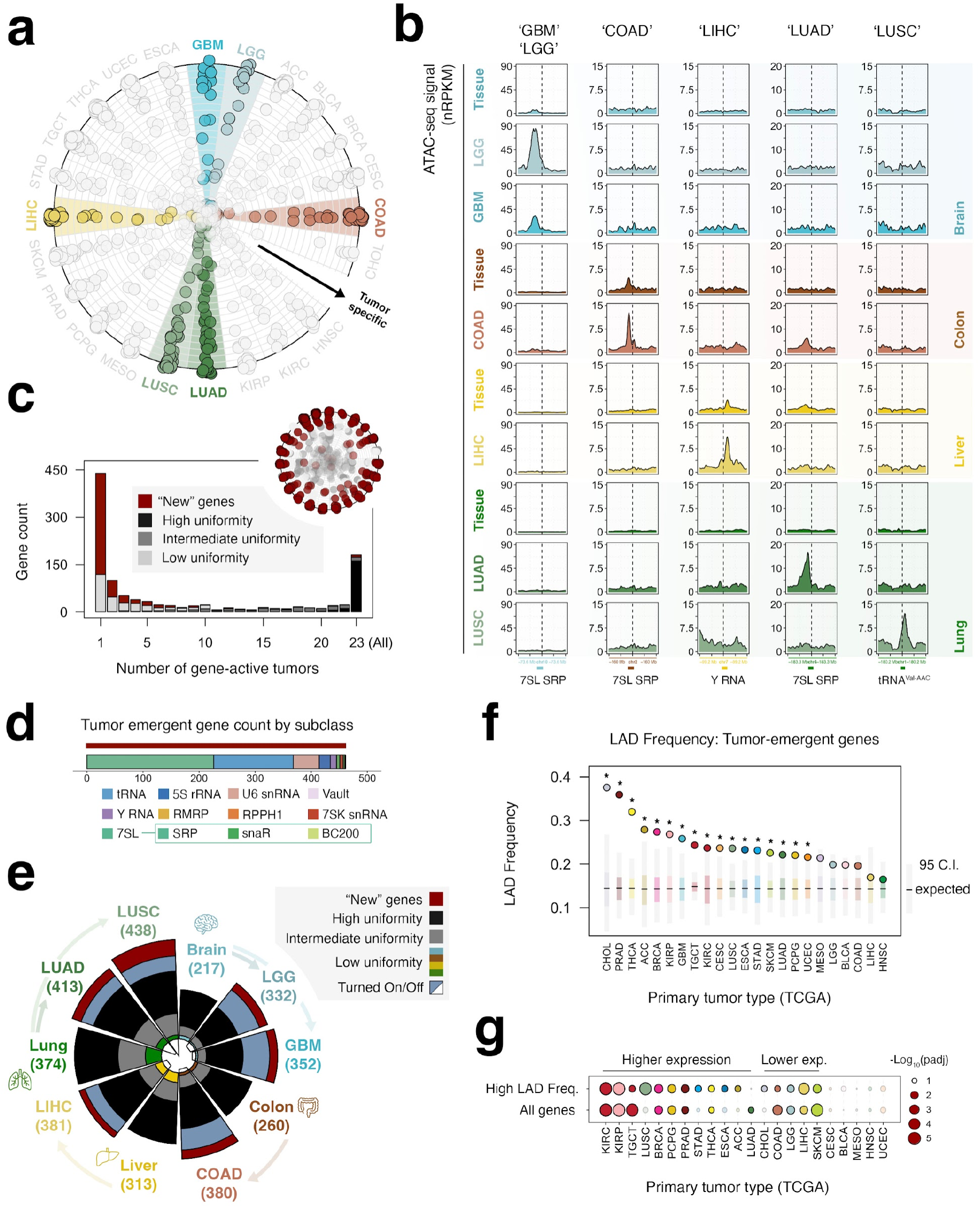
Pol III-transcribed gene patterns in cancer are dominated by tumor-specific signatures enriched within LADs. **(a)** Visualization of tumor specificity scores measured by Shannon’s entropy. Individual data points represent Pol III-transcribed genes; radially positioned genes are characterized as tumor-specific. **(b)** ATAC-seq signal pileup at specific examples of Pol III-transcribed genes defined as tumor-specific. (GBM = Glioblastoma multiforme; LGG = Brain Lower Grade Glioma; COAD = Colon adenocarcinoma; LIHC = Liver hepatocellular carcinoma; LUAD = Lung adenocarcinoma; LUSC = Lung squamous cell carcinoma). **(c)** Number of genes predicted to be active between 1 tumor subtype and all 23 tumor types. Stacked barchart denotes gene tissue uniformity (gray, black) and tumor-emergent “new” genes (red). **(d)** Composition of tumor-emergent genes by Pol III-transcribed gene subclass. **(e)** Radial barchart depicting the number of predicted Pol III-transcribed genes active in GBM, LGG, COAD, LIHC, LUAD, and LUSC, compared against relevant tissue data (brain, colon, liver, lung, respectively). **(f)** LAD (Lamina-associated domain) frequencies for tumor-emergent ‘new’ genes compared against an empirical expectation, accounting for the total new genes in each cancer subtype. **(g)** Neighborhood analysis of local expression patterns in tumors compared against randomized samples; tumors with significantly higher or lower local expression within a window size of +/-1.5 Mb (centered on tumor-specific genes) are colored; permutation p-value.

Altogether, a substantial number of the 450 cancer genes, which predominantly encode tRNA and 7SL RNA, are significantly accessible in only a single tumor type, suggesting a high degree of heterogeneity among cancer-associated Pol III patterns (Figure 5c-d). Integrating the tumor survey with the prior Pol III tissue atlas, we find that the number of predicted active genes invariably increases in tumors compared to related normal tissues (Figure 5e, Supplemental Figure 7). These patterns, such as the increased number of predicted active genes in glioblastoma multiforme or lower grade glioma compared to normal brain tissue (Figure 5e), suggest that Pol III expansion is likely a recurrent pattern in cancers.

Given that tissue-specific genes derive from regions with higher LAD frequency, we hypothesize that Pol III expansion may evolve through opportunistic activation of genes that inappropriately subcompartmentalize away from the nuclear lamina, either through context-related or entirely stochastic mechanisms. Lamin DamID analyses do confirm significantly higher LAD frequency for tumor-specific genes compared to empirical null distributions, particularly for genes linked to cholangiocarcinoma (CHOL), prostate adenocarcinoma (PRAD), and thyroid carcinoma (THCA, Figure 5e). Taken further, we additionally explored whether tumor-emergent Pol III signatures are related to changes in local mRNA expression using a gene-neighborhood analysis framework. By comparing local cancer-specific expression levels against a pan-cancer derived expectation, we find that 12 out of 22 tumor types have statistically significant deviation – both higher and lower – suggesting tumor-specific re-wiring of local expression patterns in these contexts (Figure 5g). When restricting this analysis to gene neighborhoods associated with high LAD frequency, this number increases to 16 out of 22 tumor types (Figure 5g), further indicating that tumor-specific Pol III patterns may be closely related to dynamic gene-level changes at chromatin that is ostensibly silenced at the nuclear lamina under normal conditions.

### Lamin overexpression enhances Pol III activity signatures at tumor-specific genes

Alterations in the nuclear lamina can contribute to cancer phenotypes by promoting abnormal nuclear morphology, genome instability, and dysregulation of gene expression^58^. Aberrant lamin protein levels, for example, can modulate nuclear stiffness and reprogram chromatin organization and gene expression in ways that favor tumor progression. Here, we note that high expression of Lamin A/C (*LMNA*), Lamin B1 (*LMNB1*), and Lamin B2 (*LMNB2*) are negative prognostic factors in pan-cancer survival analyses and, collectively, among the most significant negative factors compared to all HGNC gene groups (Figure 6a)^59–62^. We therefore explored the effects of *LMNA, LMNB1*, and *LMNB2* overexpression in HEK293 human embryonic kidney cells, given that kidney renal clear cell carcinoma (KIRC) includes emergent Pol III patterns enriched within high frequency LADs with neighborhood-level signatures (Figures 6b; 5f-g).

**Figure 6.**
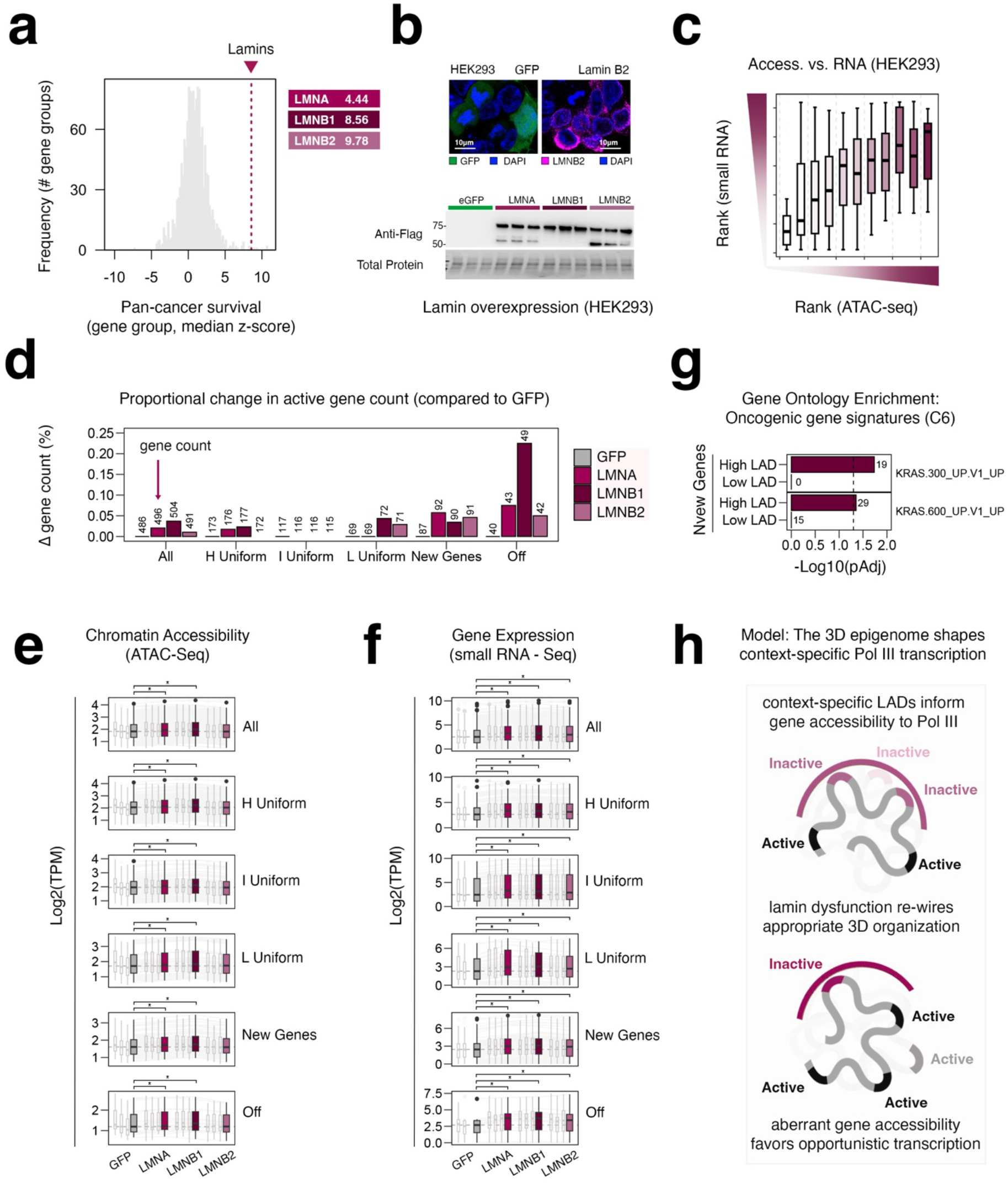
Aberrant lamin expression, an unfavorable hallmark in cancer, leads to expansion of Pol III-transcribed gene targets. **(a)** Distribution of median pan-cancer survival z-scores across 1,498 unique HGNC (Hugo Gene Name Consortium) gene groups. High z-scores are linked with unfavorable outcomes in cancer (tcga-survival.com). **(b)** Immunoblot analysis of lamin protein abundance in HEK293 cells following overexpression of either GFP (control), lamin A (*LMNA*), lamin B1 (*LMNB1*), or lamin B2 (*LMNB2*). **(c)** Integrative analysis of gene accessibility (ATAC-seq) and ncRNA abundance (small RNA-seq) in HEK293 cells. **(d)** Number of genes predicted to be active in HEK293 overexpression experiments, stratified by subcategories defined in the multi-tissue and tumor atlases. H (High), I (Intermediate), L (Low), New (tumor active), Off (active in HEK293, inactive in tissues; tumors). Y-axis depicts the proportional change in total active genes. **(e-f)** Distribution of gene accessibility (e) and RNA abundance (f) at Pol III-transcribed genes, stratified by gene groups defined in panel d. TPM = transcript per million. **(g)** Gene ontology enrichment analysis for oncogenic gene signatures (MSigDB) at New genes located within intervals of high or low LAD frequency. **(h)** Model illustration: Lamin-associated domains and cancer-related lamin dysfunction are linked with tissue- and tumor-specific Pol III patterns.

Integration of ATAC- and small RNA-seq in HEK293 confirms that gene accessibility and RNA abundance are positively linked in these contexts (Figures 6c). By further applying our tissue- and tumor-accessibility scoring framework to predict gene-level activity in HEK293, we find that *LMNA, LMNB1*, or *LMNB2* overexpression each lead to an increase in the number of significantly accessible Pol III-transcribed genes (Figures 6d). Of note, these expansions are most prominent in cancer-emergent “new” genes, as well as in other gene loci not captured in either tissue- or tumor atlases (Figures 6d). These patterns are consistent with the significantly increased accessibility observed at all classes of Pol III transcribed genes (*LMNA* and *LMNB1*), mirrored further by significant increases in small RNA abundance (*LMNA, LMNB1*, or *LMNB2*, Figures 6e-f), suggesting perturbation of nuclear lamin levels is sufficient to induce alterations in the Pol III transcription landscape.

## Discussion

The multi-dimensional nature of Pol III transcription has remained poorly understood due to a variety of unique technical challenges and relatively limited exploration. The tissue- and tumor-specific atlases reported in this study address current deficiencies using a genomic inference that, although restricted to predicting a binary gene state, provides valuable insight on the overall topology of Pol III transcription in various contexts. Principle among these insights are the discoveries of variable Pol III expansion in tissues and tumors, a unique catalog of universal and near-universal Pol III-transcribed genes, and a novel connection between tissue-specific and tumor-emergent Pol III patterns and chromatin subcompartmentalization at the nuclear lamina. Recognizing the interplay of Pol III activity and subnuclear 3D chromatin organization is congruent with the well-established role of LADs in tissue-specific gene regulation^63,64^.

Mechanistically, we show that lamin overexpression is generally sufficient to drive analogous expansions of active Pol III-transcribed genes in a specific cellular context. While such increases are most pronounced at genes classified as tumor-emergent (“new”), we note that the magnitude of expansion is limited in nature, with an overall increase in ~ 10-20 genes (Figure 6d). We speculate that acute increases in lamin protein abundance and downstream effects on 3D organization may allow for opportunistic transcription, providing further opportunities for clonal selection in favor of changes that promote growth within heterogeneous cell populations. When considering the overall landscape of tumor-emergent Pol III genes with high LAD frequency, we find a proximal enrichment for KRAS-sensitive genes (Figure 6g). KRAS signatures are particularly notable here given the high prevalence of KRAS involvement across a broad spectrum of cancer types, suggesting Pol III gene-dynamics may be physically linked to gene neighborhoods that are recurrently affected by KRAS dysfunction^65^.

In summary, our study points to the nuclear lamina as a major player in defining the Pol III transcriptome, such that context-specific LADs actively shape the cellular small RNA landscape (Figure 6h). LAD organization itself likely functions as a large-scale tapestry, with additional gene regulatory layers (e.g. specific chromatin and transcription factors) contributing to more nuanced Pol III control. It is important to note that not all inactive genes are organized within LADs in any cell type, and thus other mechanisms are indeed required and responsible for tissue-specific Pol III patterns as well. Nevertheless, the biased expansion of Pol III in cancer (rather than contraction) indicates that loss of gene silencing, through LAD reorganization and/or other mechanisms, is the primary nature of Pol III transcription dysregulation during tumorigenesis.

### Limitations of this study

Both tissue- and tumor atlases rely on chromatin accessibility as a proxy for Pol III transcription. While this approach cannot independently distinguish sites with productive from nonproductive transcription, our strategy addresses major technical obstacles that have limited investigations to date, including challenges related to RNA sequencing and mapping. Our uniform gene accessibility framework is sufficient to capture dominant tissue and tumor patterns, though fine-scale resolution naturally remains unresolved. Additionally, the dominant patterns reported in the present study are restricted to currently available contexts, which passed stringent quality control filters. Future investigation of chromatin accessibility in expanded tissue and tumor contexts may allow for reexamination of Pol III ubiquity and specificity.

While our findings support a role for the nuclear lamina in shaping Pol III patterns, including evidence that lamin overexpression expands the repertoire of Pol III-transcribed genes – particularly at tumor-emergent loci – the precise molecular mechanisms underlying these changes remain unresolved. Lamins may influence Pol III through multiple interconnected processes, be it via 3D chromatin architecture, nuclear stiffness, DNA damage responses, etc. Though this complexity presents a challenge in pinpointing how lamin perturbations directly contribute to Pol III dynamics, the link between lamin dysregulation and Pol III expansion highlights a novel mechanism by which gene subcompartmentalization may additionally contribute to disease progression and cancer heterogeneity.

## Materials and Methods

### Cell lines and culture conditions

HEK293 (ATCC CRL-1573) were cultured in complete Dulbecco’s Modified Eagle’s Medium (DMEM) (Corning) supplemented with 10% fetal bovine serum (Corning) and 1% Penicillin-Streptomycin. Cells were maintained in 10-cm dishes by seeding 1 × 10^6^ cells with complete medium every 2-3 days. Cells were maintained in a humidified atmosphere at 37 C with 5% CO2. H1 human embryonic stem cells (H1-hESCs) were obtained from WiCell (Wisconsin, USA). H1-hESCs were grown in Matrigel (Corning, NY, USA)-coated 12-well plates in Essential 8™ Medium (Thermo Fisher Scientific, MA, USA) at 37°C in 5% CO_2_ in compressed air and high humidity. Cardiomyocyte differentiation relied on previously established protocol that generates beating cardiomyocytes derived from H1-hESCs^66^. To further increase cardiomyocyte purity, the differentiated cells were subjected to subsequent glucose starvation using non-glucose-supplemented RPMI/B27 medium for 3× (2 days per time) to decrease noncardiomyocyte cells because cardiomyocytes are more tolerant to glucose starvation. Following a 30-day differentiation protocol, cells were collected using Accutase (Thermo Fisher Scientific).

### Plasmids and cell transfection

pcDNA3.1+/C-(K)DYK Plasmids expressing eGFP, LMNA (Catalog#OHu27661D), LMNB1 (Catalog#OHu20925D) and LMNB2 (Catalog#OHu21383D) were obtained from GenScript. HEK293 cells (1 x 10^5 cells/cm2) were seeded into 12-well Cell-Culture Treated Multidishes (Thermo Scientific) and incubated overnight. Plasmids were transfected using Lipofectamine 3000 (Invitrogen) according to the manufacturer’s protocol. Cells were collected 48 hours post-transfection for subsequent analysis.

### Immunoblotting

Cell pellets were collected after transfection and washed once with PBS before lysis with cell disruption buffer (Thermo Scientific) for 5 minutes on ice, followed by centrifugation at 14000 g for 5 minutes at 4 C. The lysates were separated on a 4 - 20% Mini-PROTEAN TGX Stain-Free Protein Gels (BIO-RAD and transferred to a polyvinylidene difluoride membrane. Transferred membranes were blocked with 5% blotting grade blocker in PBST and followed by incubation with the primary antibody at 4 C overnight. Membranes were washed 3×5min with PBST and incubated with the secondary antibody for 1 hr at room temperature, followed by 3 washes with TBST. Proteins were visualized using SuperSignal WestFemto (ThermoScientific) with CehmiDoc Touch Imaging System (BIO-RAD)

### Immunofluorescence

HEK293 cells were cultured on PDL-coated 8-well chamber slides (ibidi) and washed once with PBS (Corning). Samples were fixed in 4% paraformaldehyde (Thermo Scientific) for 15 min at room temperature, followed by two washes in PBS (5 min each). For permeabilization, cells were incubated with 0.5% Triton X-100 for 10 min on ice, then washed 3 times in PBS (10 min each). Non-specific binding was blocked by incubating samples with 1% BSA (MP Biomedicals) in PBS for 30 min at room temperature. Samples were incubated with LaminB2 primary antibody (Cell Signaling Technology E1S1Q, 1:100) at 4 °C overnight, followed by three PBS washes (10 min each). Secondary antibody was applied at 1:500 dilution for 1h at room temperature in the dark. After three additional PBS washes (10 min each, in the dark), nuclei were counterstained with 5 µg/ml DAPI. Wells were mounted using antifade mounting medium and proceeded with imaging on a ZEISS LSM 900 confocal using a 60× oil objective.

### DNA Library preparation and sequencing

For ATAC-Seq, 100,000 HEK293 cells were collected following overexpression of eGFP, LMNA, LMNB1, or LMNB2. Libraries were prepared using the ATAC-Seq Kit (Active Motif) according to the manufacturer’s protocol. For small RNA-Seq, cells from the same overexpression conditions (eGFP, LMNA, LMNB1, or LMNB2) were collected and small RNA extracted using the mirVana PARIS Kit following the manufacturer’s instructions (Thermo Fisher Scientific). ATAC and Small RNA libraries were prepared and sequenced on the NovaSeq X Plus (Illumina) platform for 150 and 100 cycles respectively. All sequencing was performed at the University of Illinois, Roy J. Carver Biotechnology Center.

### POLR3A ChIP-seq

H1 stem cells and H1-derived cardiomyocytes were harvested (~10 million cells per ChIP experiment) and resuspended in growth media at 1 × 106 cells/mL and cross-linked with rotation at room temperature in 1% formaldehyde for 10 min. Cross-linking was quenched with the addition of 200 mM glycine and an additional 5 min of rotation at room temperature. Cross-linked cells were then spun down and resuspended in 1× RIPA lysis buffer, followed by chromatin shearing via sonication (3 cycles using a Branson sonicator: 30 s on, 60 s off; 20 additional cycles on a Bioruptor sonicator: 30 s on, 30 s off). Individual ChIP experiments were performed on pre-cleared chromatin using antibody-coupled ChIP grade Protein G magnetic beads (Cell Signaling Technology). POLR3A antibody was obtained from ThermoFisher Scientific (PA5-58170). 5 ug of antibody per ChIP was coupled to 18 uL of beads and rotated overnight with sheared chromatin at 4°C. Beads were then washed 5× in ChIP wash buffer (Santa Cruz), 1× in TE, and chromatin eluted in TE + 1% SDS. Cross-linking was then reversed by incubation at 65°C overnight, followed by digestion of RNA (30 min RNase incubation at 37°C) and digestion of protein (30 min proteinase K incubation at 45°C). ChIP DNA was then purified on a minElute column (Qiagen), followed by DNA library preparation (NEBNext Ultra II DNA Library Prep Kit for Illumina) and size selection of 350-550 bp fragments via gel extraction (Qiagen).

### ATAC-Seq data processing

We integrated differentiated tissues ATAC-Seq data from the Encyclopedia of DNA Elements (ENCODE) and the Gene Expression Omnibus (GEO) series GSE164942 (Part of Liver data) and GSE96949 (Brain data) and primary solid tumors from the Cancer Genome Atlas (TCGA), collecting more than 1000 different datasets. The processing of ATAC-Seq data was carried out by following and adapting established protocols. Reads were first trimmed using TrimGalore and then mapped to the genome (GRCh38) using BowTie2. Counts were extracted over a comprehensive RNAcentral database annotation of noncoding RNAs Using BEDTools. For each gene, a total of

150 nucleotides were added to both the upstream and downstream regions to improve mapability. Additionally, counts were extracted from the regions upstream and downstream of the gene of interest. These regions were 500 bp, 5000 bp, and 50000 bp in size with a buffer region of 300 bp allowed between the genes and the additional mapped regions. Technical replicates were further aggregated by adding the counts in each one of the respective samples except for the data obtained from GSE96949 where counts were aggregated across technical and biological replicates. The rationale behind this is related to the low sequencing depth observed in these experiments compared to other data collected. ATAC-Seq files from Lamin overexpression were processed similarly, with subsequent TPM normalization.

### ChIP-Seq data processing

We integrated ChIP-Seq data from ENCODE, a total of 47 experiments, focusing on six specific Pol III subunits: POLR3A, POLR3B, POLR3C, POLR3D, POLR3E and POLR3G. Reads were trimmed using TrimGalore and then mapped to the genome (GRCh38) using BowTie2. Similarly to ATAC-Seq data, we extracted counts over the same RNAcentral database annotation of noncoding RNAs. We aggregated all the reads found within the gene coordinates for each respective Pol III subunit across all experiments. After aggregation, we normalized the data and calculated the upper quantile for each unique subunit across all experiments. This statistical measure helped identify the higher concentration of Pol III occupancy within the genome, serving as a benchmark for assessing significant Pol III activity. Furthermore, to determine a global estimate of Pol III occupancy, we calculated the median value across all six Pol III subunits. ChIP-Seq data for POLR3A subunit from hiPSC and differentiated neurons were obtained from the GEO series GSE227928. POLR3A ChIP Seq data was processed in the same way as described. Raster plots were obtained by merging BAM files and converting them into a BigWig file and subsequently RPKM normalized using Deeptools.

### Small RNA-Seq data processing

RNA reads were trimmed using TrimGalore and eventually mapped to the genome using (GRCh38) BowTie2. Counts were extracted over the same RNAcentral database annotation of noncoding RNAs and subsequently TPM normalized.

### Gene accessibility scoring

Gene accessibility scoring across the different gene annotations was determined by assuming that read alignment on the genome follows a Poisson distribution. Briefly, for each gene, the expected signal within 1 kb 10 kb 100 kb and across the genome was calculated. Then the maximum value of the signal is estimated to be lambda (*λ*). The probability density function (PDF) used is given by *P*(*Xi* = *K*) = *λ*^k * e ^-*λ*/k!, where *P*(*Xi* = *K*) represents the probability of observing K aligned reads for the gene i given (*λ*), and 1 - *P*(*Xi* = *K*) represents the p-value tied to each bin. P-values were adjusted globally using the Benjamini-Hochberg false discovery rate method. (https://github.com/VanBortleLab/Pol_III_tissue_tumor_atlas)

### Sequencing depth analysis

To calculate optimal sequencing depth for all the experiments, each sample was downscaled by factors ranging from 0.05 to 0.9, with counts rounded down to the nearest whole number. Subsequently for each factor, gene accessibility scores were calculated as described previously. The total number of genes deemed significant before downscaling served as a baseline for comparison with the number of significant genes post-downscaling. A non linear model was applied to estimate the optimal sequencing depth - Y = (Ymax*SeqDepth) / (Kte + SeqDepth) - where Y is the proportion of significant genes, Ymax is the maximum proportion of significant genes, Kte is the function constant that serves as a change rate in the proportion of significant genes and SeqDepth is the sequencing depth. This equation can be linearized by taking the reciprocal to estimate Ymax and Kte. By relying in this method, combined with visual inspection, we downscaled all samples to a total of 250,000,000 reads. Samples with fewer reads before downscaling or those retaining less than 90% of the total gene count post-downscaling were omitted from the analysis, being categorized as noise. No sample was upscaled.

### Cumulative signal plot generation

To generate raster plots for all tissue types and cancer samples, BAM files were merged and converted into a BigWig file. These files were aligned to the genome, downscaled by a specified factor, and normalized using Deeptols version 3.5.6. Normalization was performed with the following command line: bamCoverage —binsize 20 – scaleFactor –smoothLength 60 –normalizeUsing RPKM. Afterward, BW files from samples belonging to the same tissue types using Deeptools, with the command bigwigAverage –binSize 20.

### RNA Polymerase III Gene Ubiquity Classification

To score tissue heterogeneity within the Pol III transcriptome, we used Shannon’s entropy. Samples were grouped into their respective tissue, and three distinct entropy values were computed. Specifically, 134 samples were consolidated into 19 tissue samples, with samples not fitting any category labeled as ‘other. First, a binary entropy based on the gene’s binary on/off status was calculated. Second, a p-value entropy was derived from the negative log p-value to assess gene significance. Lastly, count entropy was determined by the proportion of samples within a tissue where a gene was considered significant. For each gene, a relative score (p(x)) and entropy (H) were calculated as follows. p(x) = score/Σscore and H = −Σp(x)*log2(p(x)). The maximum value for H is equal to the log2(N) and N is the total number of samples. Three different groups were defined for tissues according to their entropy values considering high, intermediate, and low uniformity genes have H > log2(14), H > log2(4), and H < log2(4) respectively. All these genes were considered as ON.

### Tissue and Cancer dominance visualization

Gene accessibility scores were binarized across all the experiments and then median binary score was calculated for the experiments belonging to a specific tissue or cancer. Dominance across multiple variables is calculated based on their entropy level. Observations concentrated in the center of the circle are significantly accessible across all variables and as an observation starts approaching the perimeter of the circle it becomes more accessible. Concentric circles within the plot represent the degree of specificity accounting for the number of variables an observation is shared, being the outermost a single variable. plot_circle() calculates the entropy and plots specificity of observations (https://github.com/VanBortleLab/dominatR).

### tRNA sequence analysis prediction

We selected tRNA genes and conducted discriminative motif discovery using sequences of 300 bp centered on the transcription start sites (TSS) of all tRNAs, utilizing the MEME-Suite. For the motif discovery, we employed STREME with the following parameters: --dna --minw 8 --maxw 15 --order 5 --thresh 0.5 --align left --p. After identifying specific motifs, we scanned these against all tRNA sequences using Fimo. We then used the scores from Fimo to predict gene activity states (ON vs. OFF) and expression levels (High-Intermediate Frequency vs. Low Frequency) employing two classifiers: random forest and penalized logistic regression. Random forest outperformed penalized logistic regression, leading us to conduct subsequent analyses with this model. For assessing variable importance, we utilized the R package Boruta, which is based on random forest. To identify potential motifs that matched our enriched sequences, we used Tomtom, integrating sequences from JASPAR2022_CORE_ vertebrates, HOCOMOCOv11_full_Human, and previously reported A and B boxes.

### Epigenetic features analysis

We integrated data from ChipAtlas to analyze histone markers, chromatin accessibility, transcription factors, and DNA methylation and from 4DNucleome to analyze Lamina Associated Domains (LAD). We processed all the bigwig files, extracting the normalized signal from a genomic region of 1 kb centered on the transcription start sites (TSS) of all Pol III genes, using a bin size of 10 bp. For histone markers, chromatin accessibility, and DNA methylation, we included experiments with over 5000 peaks, as identified by ChipAtlas. For transcription factors (TFs), we considered only those experiments that featured more than 40 datasets. For LADs we retrieved Dam-Id and pA-Dam sequencing experiments subsetting for LMNB1, LMNB2 and LMNA/C. Additionally, for each condition, we calculated the median and the upper quartile and derived two global scores for each gene by summing or extracting the maximum signal across the 1 kb region. In total, we had 4 different dataframes:

- Experiment Score: Median & Gene Signal: Max
- Experiment Score: Upquar & Gene Signal: Max
- Experiment Score: Median & Gene Signal: Sum
- Experiment Score: Upquar & Gene Signal: Sum

For each dataframe, we determined variable importance and accuracy scores using a random forest classifier to predict gene activity states (ON vs. OFF) and expression levels (High-Intermediate Uniformity vs. Low Uniformity). We generated 1000 different Training-Test splits where 500 kept the structure of the response variable and the other 500 shuffled the response variable. We summarized the scores of each analysis across each dataframe upon normalization by calculating the mean score.

### Lamina Associated Domains Frequency and nuclear positioning analysis

The definition of Lamin Associated Domains (LADs) involved leveraging publicly available datasets from the 4D Nucleome Data Portal. We focused on BED files from DNA adenine methyltransferase identification sequence (DamID-Seq) and proteinA-DamID (pA-DamID) experiments, specifically targeting LMNB2, LMNB1, and LaminA/C proteins. These files contain the coordinates for the LADs. Biological experiments produced multiple BED files that define LADs at various bin resolutions. We consolidated entries from each unique experiment by considering the frequency of LAD calls across different bin resolutions. The intra-experimental LAD calls were determined using BEDTools multiintersect. For each experiment, we limited LAD calls to those coordinates with the highest LAD frequency. In this context, LAD frequency can be defined as the overlap of specific coordinates across all experiments, meaning that coordinates with high frequencies are observed across all the different experiments. This comprehensive analysis spanned 111 unique experiments, allowing for a global overview of LAD distribution within the genome. To facilitate wider access and further research, we have established a repository on GitHub, serving as a repository for the data and findings (https://github.com/VanBortleLab/LADFreq).

The primary objective of determining LAD frequencies was to examine the enrichment of previously defined gene groups within these specific genomic regions. To assess the extent of overlap between our identified LADs and the genes of interest, we employed the BEDTools intersect function with settings ‘bedtools intersect -a -b -wao’. This approach enabled us to quantify the degree of LAD frequency overlap for each gene in each group. The strength of LAD frequencies overlap per group was determined by comparing these observations against a non-parametric distribution derived from a permutation test (n = 100,000), while controlling for the number of genes per group.

Nuclear positioning information was obtained from the genomic annotation BED files provided in the GEO series GSE135582. These coordinates were originally annotated to the GRCh37 (hg19) genome assembly. Prior to overlap analysis, all coordinates were converted to GRCh38 (hg38) using the UCSC LiftOver tool (genome.ucsc.edu). Following coordinate conversion, overlaps between GPS-Seq–scored loci and the genes of interest were assessed using the same strategy described for Lamin-associated domain (LAD) frequency analyses.

### Neighborhood analysis of local expression

Neighborhoods were defined as protein-coding genes located in proximity to Pol III–transcribed loci associated with emergent *New Genes* in tumors. To establish the set of protein-coding neighbors, we used RNA-Seq data matched to the ATAC-Seq samples analyzed in this study. RNA-Seq expression dataframes were obtained for matched samples, and only protein-coding genes were considered. For each tumor type, genes in the lowest expression quartile were excluded as lowly expressed. For each Pol III–transcribed locus, the 10 closest protein-coding genes were identified using BEDTools closest (-d -t first -io)^67^. Protein-coding neighbors located farther than 1.5 Mb upstream or downstream of the Pol III transcription start site were excluded; furthermore, only neighborhoods containing ≥3 protein-coding genes after filtering were retained.

For each neighborhood, expression was summarized by computing the mean across all nearby genes. Neighborhood expression datasets were generated for each tumor type by collapsing matched RNA-Seq data for the respective neighborhoods. Tumor-specific neighborhood activity was defined as the expression of all neighborhoods associated with *New Genes* active in that tumor, and the same set of neighborhoods was extracted across all other tumor types for comparison.

To assess whether tumor-specific neighborhoods displayed differential expression relative to other tumors, we implemented a permutation-based resampling test. For each cancer type, we compared the observed median expression of its active neighborhoods against a null distribution generated by randomly sampling expression values (100,000 iterations). For each neighborhood set, both upper- and lower-tail empirical p-values were computed and adjusted for multiple comparisons using the false discovery rate (FDR) method.

To further evaluate the impact of nuclear compartmentalization, Pol III neighborhoods with LAD overlap frequencies above 0.15 were retained. The permutation procedure described above was repeated to assess neighborhoods occurring in high LAD frequency regions.

### Functional enrichment analysis

Gene Ontology enrichment analysis was conducted using the R package cluster Profiler with genome-wide annotations for Human.org.Hs.eg.db. Overrepresentation analysis was conducted on the C6 : Oncogenic signature gene sets^68–70^. Gene sets were defined based on the neighborhoods of the *New Genes* identified in this study. Two specific sets were of particular interest: genes located in neighborhoods with low LADs and genes located in neighborhoods with high LADs. These were compared against a background universe consisting of all genes present in the neighborhoods of all *New Genes*. Statistical significance was assessed, and p-values were adjusted using the Benjamini–Hochberg method.

### Survival Analysis

Clinical signatures, which link mRNA expression with TCGA patient outcome data, were retrieved from tcga-survival.com and correspond to cancer-specific and summary z-scores calculated using Cox univariate hazard models. Gene group information was retrieved from the Hugo Gene Nomenclature Committee (HGNC) from https://www.genenames.org/download/custom/, selecting for “Gene Group Name”. For each unique gene group, the median pan-cancer survival score was calculated with further comparisons drawn from the resulting frequency distribution.

## Resource Availability

The ChIP-seq, ATAC-seq, and RNA-seq data generated for this study are available through the NCBI Gene Expression Omnibus with accession numbers GSE306220 and GSE306294.

## Acknowledgements

We thank Alvaro Hernandez, Chris Wright, Danman Zhang, and staff at the Carver Biotechnology Center for sequencing services, and administrators of the Carl R. Woese Institute for Genomic Biology (UIUC) Biocluster for computational support. We thank members of the Van Bortle lab for helpful suggestions, and Prof. Andrew Belmont’s lab for generously sharing Alexa Fluor 647 secondary antibody. This work was supported by the National Institutes of Health, National Human Genome Research Institute (NHGRI) grant R00HG010362 and funding from the Roy J. Carver Charitable Trust to K.V.B; startup funding from Clemson University and the SPARK-South Carolina Alzheimer’s Disease Research Center pilot grant to Q.L.

## Author contributions

Study design: SL, KVB. Data collection: SL, SZ, RC, KVB, YS, QL. Data analysis: SL, SZ, KVB. Data interpretation: SL, KVB. Writing: SL and KVB with comments from all authors.

## Supplemental Figures

**Supplemental Figure 1.**
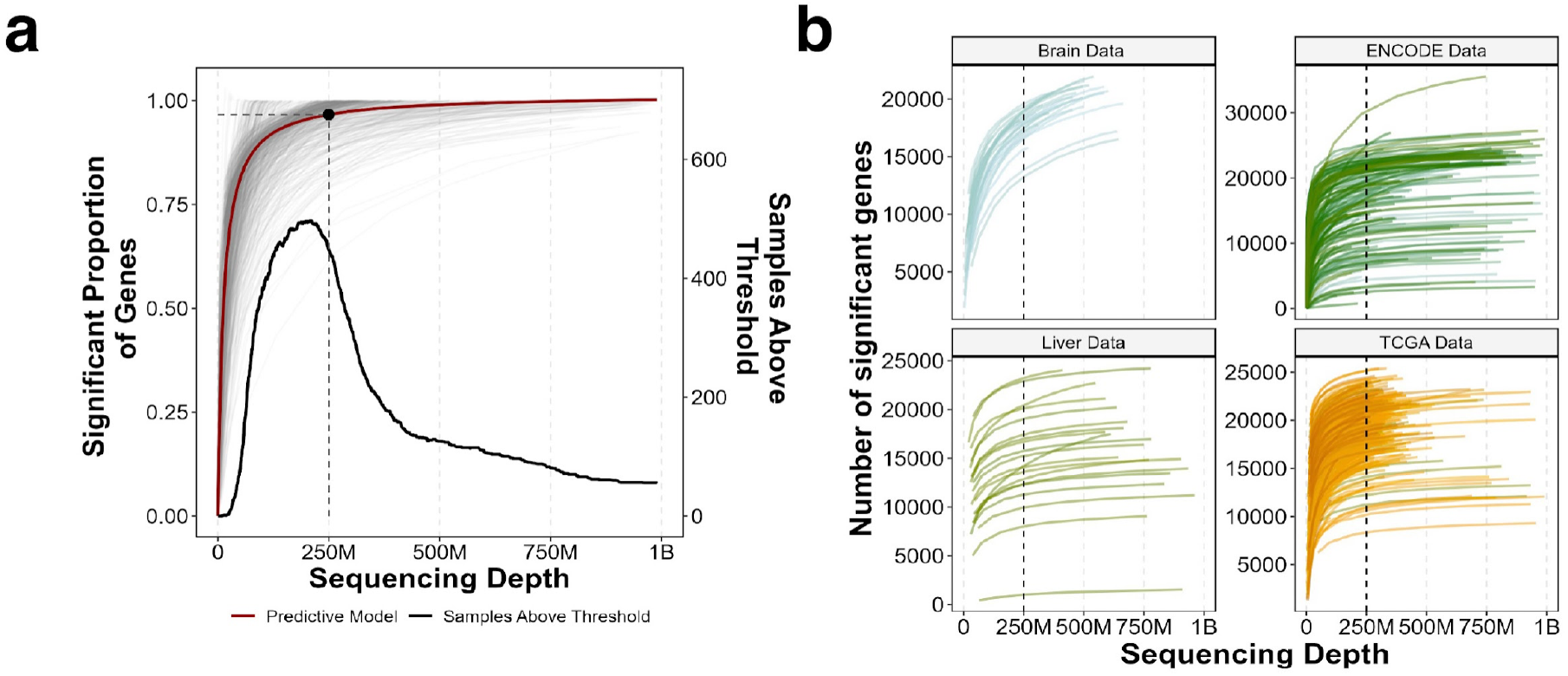
Prediction-based saturation analysis across 699 ATAC-seq datasets. **(a)** Fraction of annotated genes determined to be active at maximum sequencing depth (i.e. maximum “on” set) re-analyzed at variable levels of downscaling (individual biological ATAC study samples visualized in grey, multi-sample regression in red). A sequencing depth of 250 million reads was chosen as an arbitrary threshold based on > 95% saturation estimation, and the number of ATAC-seq datasets with sufficient depth (black line denotes the distribution of experiments at variable levels of sequencing depth; dotted line serves as a reference for 250 million reads). **(b)** Visualization of saturation curves for ATAC-seq datasets segregated by resource (TCGA = The Cancer Genome Atlas). In total, 462 ATAC-seq datasets were included based on sufficient sequencing depth (>= 250 million reads) and sufficient saturation after downscaling.

**Supplemental Figure 2.**
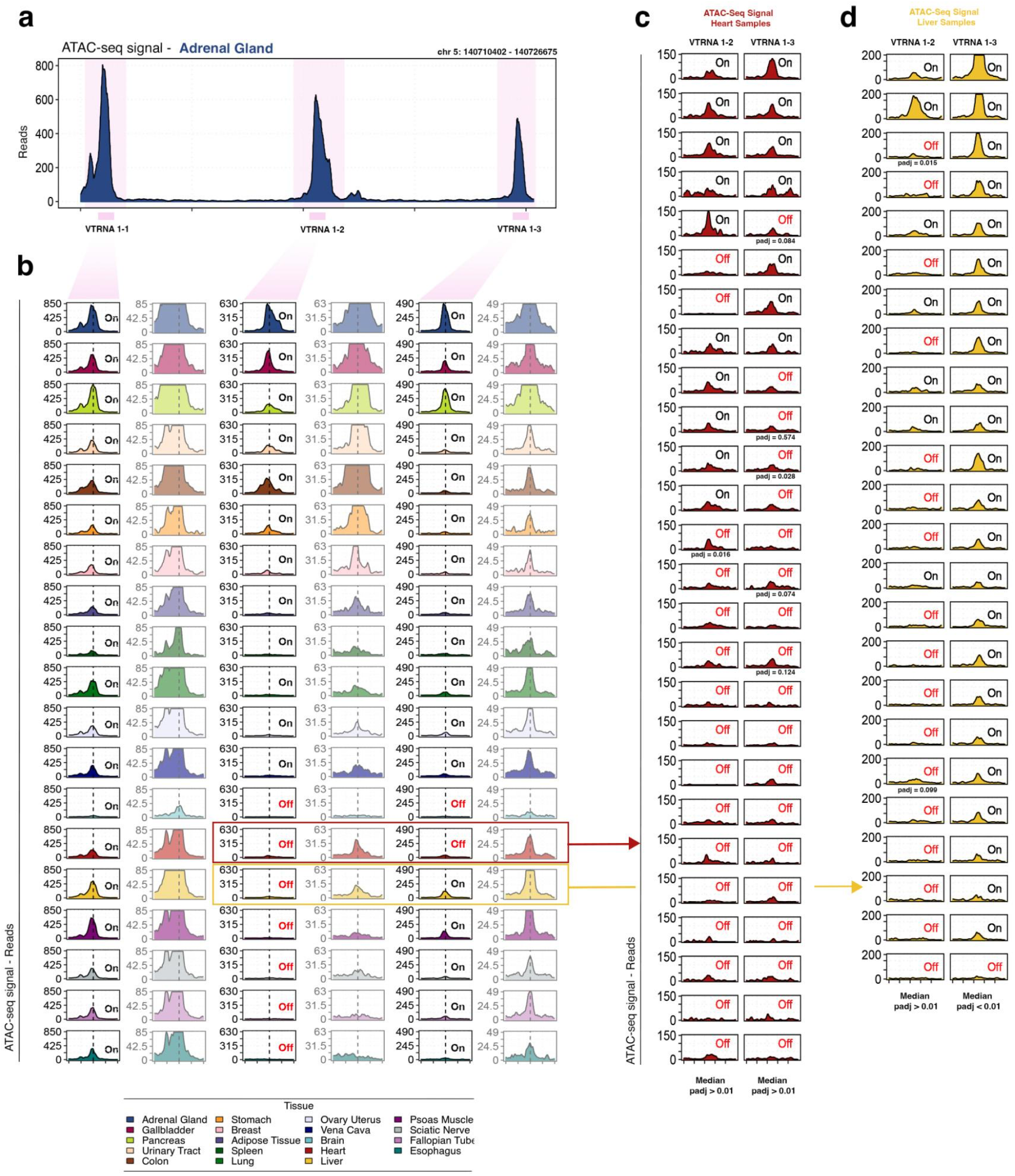
Inspection of individual vault RNA genes, which include examples of shared and tissue-restricted gene signatures. **(a)** ATAC-seq signal pileup at the vault RNA gene locus on chromosome 5, which includes vtRNA1-1, vtRNA1-2, and vtRNA1-3, in adrenal gland samples, a context predicted to be active. **(b)** ATAC-seq signal pileup at the vault RNA gene locus across all 19 tissues (tissue legend at bottom). Vault RNA gene vtRNA1-1 is predicted to be active across all 19 tissues (high uniformity, or “shared”), whereas vtRNA1-2 and vtRNA1-3 are characterized as having varied levels of context-restricted patterns (intermediate uniformity). Each gene-centered pileup is shown at uniform maximum scaling and complemented with reduced maximum scaling for clarity. **(c-d)** Deeper inspection of ATAC-seq signal pileup at all experiments underlying heart (c, red) and liver (d, yellow), which exemplify sites with varied accessibility at the boundary of being called “on” or “off”.

**Supplemental Figure 3.**
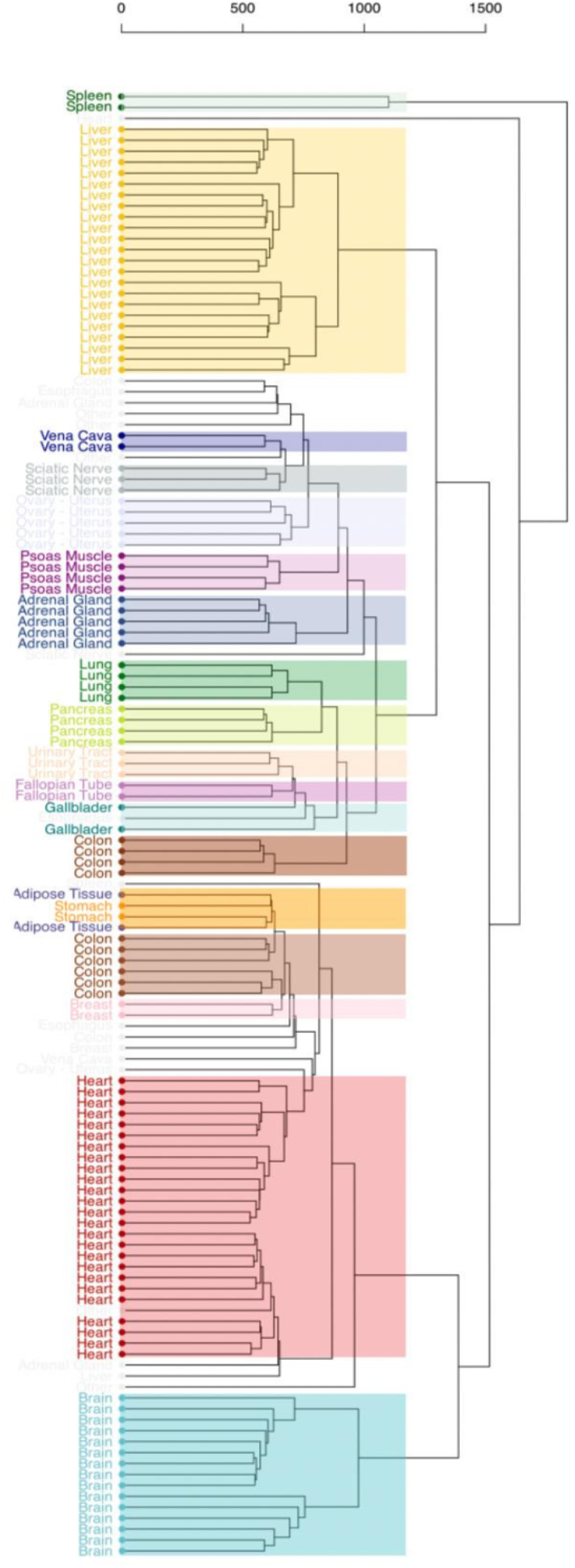
Hierarchical clustering of 134 ATAC-seq datasets spanning 19 human tissues subtypes based on Pol III gene activity signatures. Individual experiments are categorically colored by human tissue, as shown in Figure panel 2a, but with emphasis on all 19 distinct human tissue types.

**Supplemental Figure 4.**
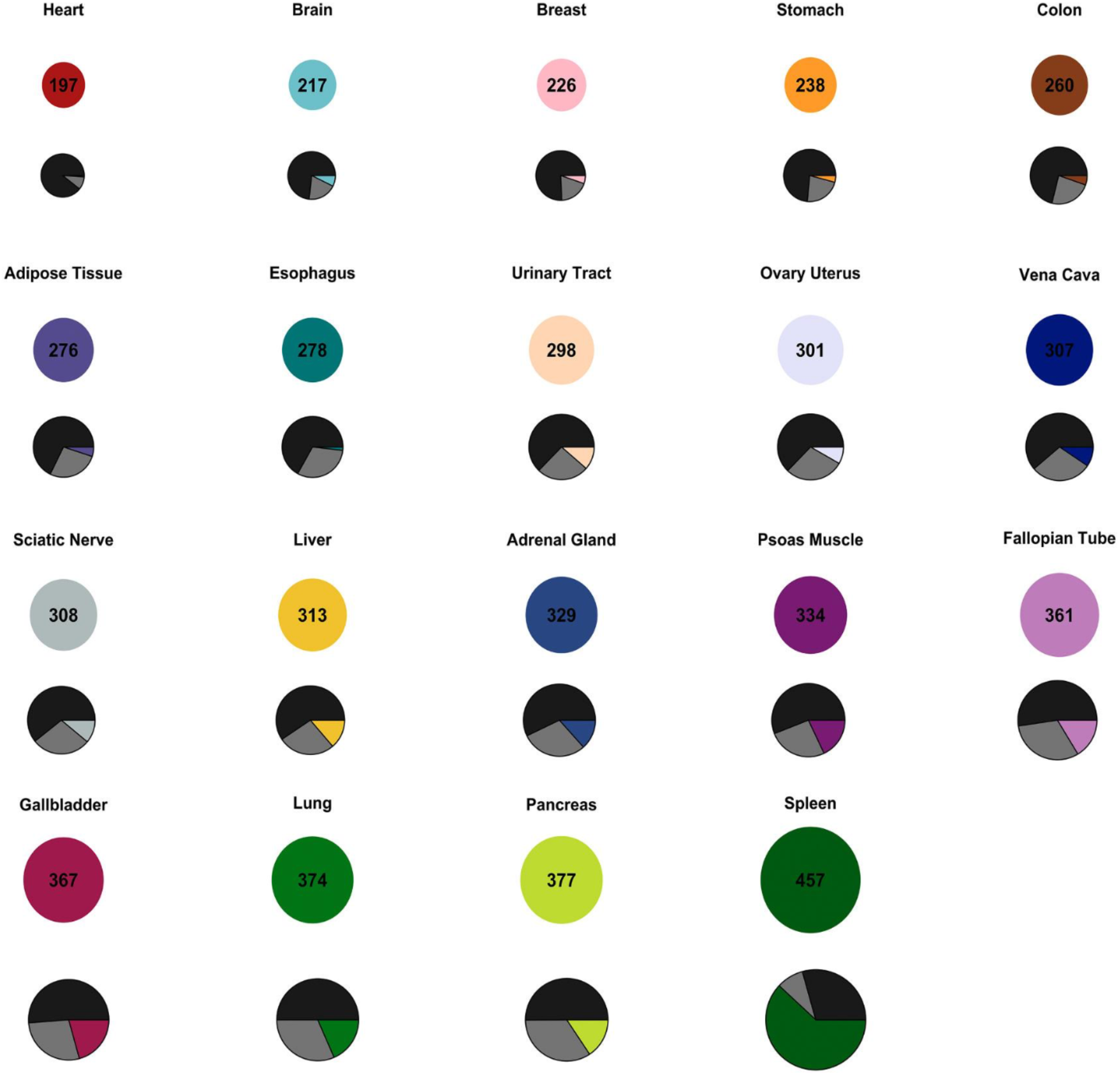
Overview of the variable breadth and number of tissue-specific Pol III-transcribed gene features defined across all 19 human tissue subtypes (related to figure panel 2b). Pie charts denote the number of predicted active “ON” genes in each tissue, and the fractional composition of high uniformity (black), intermediate uniformity (grey), and low uniformity (tissue-specific color) Pol III-transcribed genes in each tissue.

**Supplemental Figure 5.**
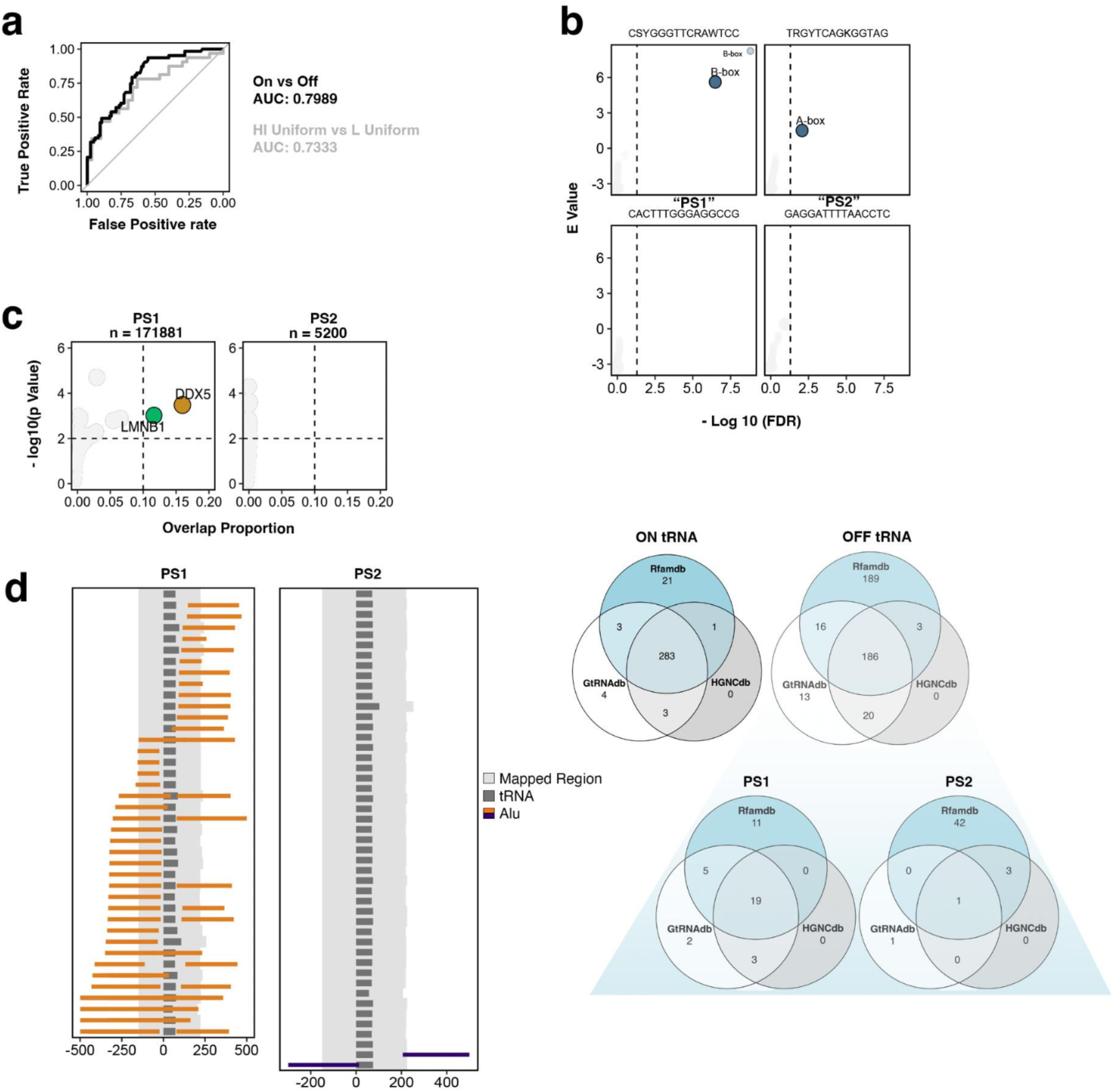
Sequence feature analysis at Pol III-transcribed tRNA genes (related to Figure 3). **(a)** Classification model performance for defining active vs. inactive tRNA genes (black) and high vs. low uniformity (grey); AUC = Area Under Curve. **(b)** Statistical output from motif comparison (TOMTOM) analysis for predictive sequences, including matches to A-box and B-box (top) and the absence of matching motifs for predictive sequences 1 and 2 (PS1 and PS2, respectively). E = enrichment value. **(c)** Statistical output following chromatin and transcription factor (TF) overlap analysis at PS1 and PS2. ChIP-seq data obtained from ChIPatlas. **(d)** Positional mapping of Alu sequences (orange; purple) with respect to tRNA gene loci annotated with PS1 or PS2. Venn diagrams illustrate the number of tRNA genes defined as being “ON”, “OFF”, and “OFF + PS1” or “OFF + PS2”, and the underlying gene annotation source.

**Supplemental Figure 6.**
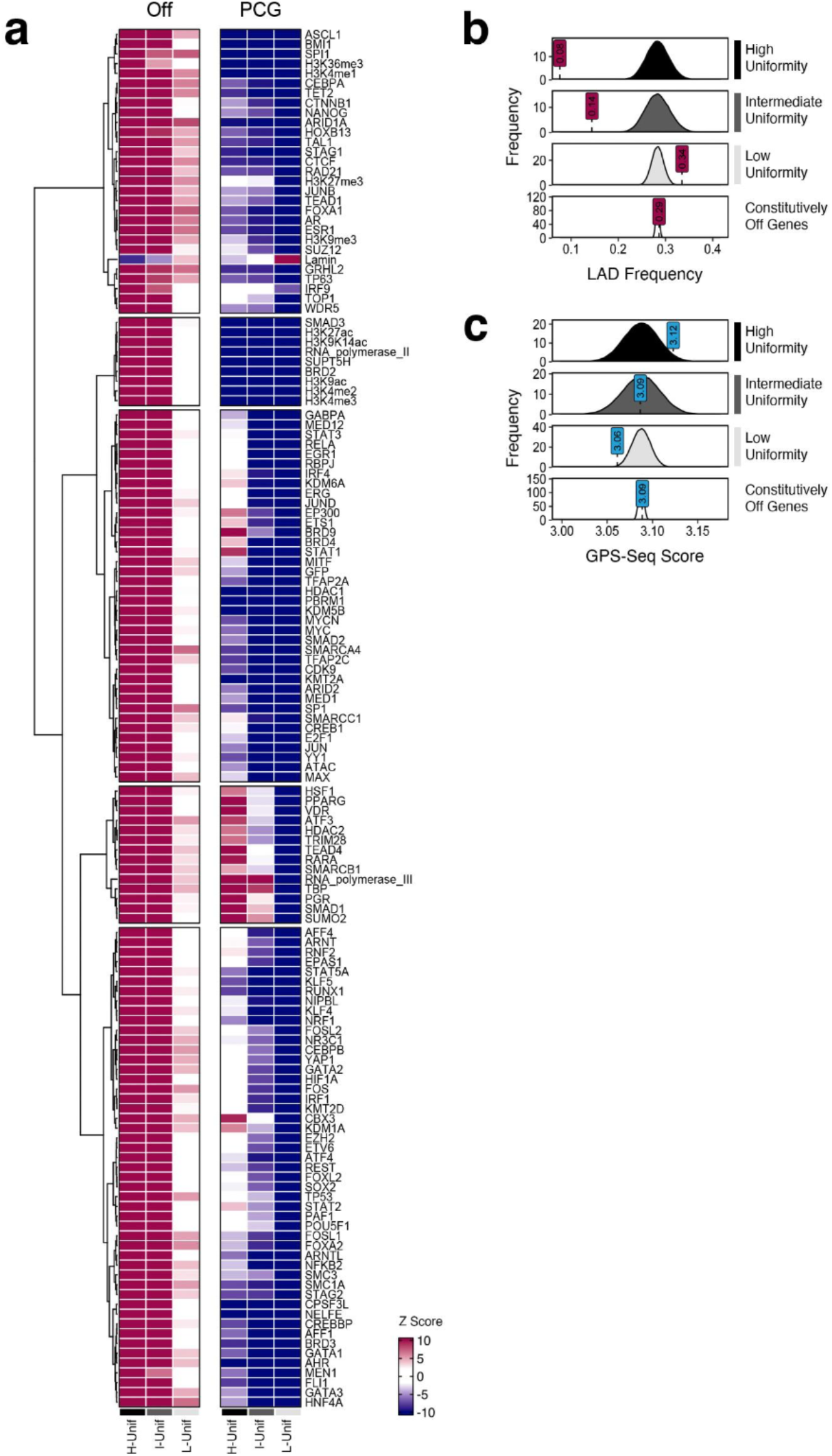
Epigenome analysis at Pol III-transcribed genes stratified by tissue-ubiquity. **(a)** Heatmap of feature enrichment z-scores at high, intermediate, and low uniformity genes. H = High uniformity; I = Intermediate uniformity; L = Low uniformity. Off (all annotated, inactive Pol III-transcribed genes) and PCG (all protein-coding genes) refers to the underlying null distribution. **(b)** Observation and empirical null distributions for average LAD frequencies spanning high, intermediate, and low uniformity genes, using the inactive Pol III-transcribed genes as null distribution. **(c)** Analogous analysis corresponding to radial subnuclear positions mapped by GPS-seq.

**Supplemental Figure 7.**
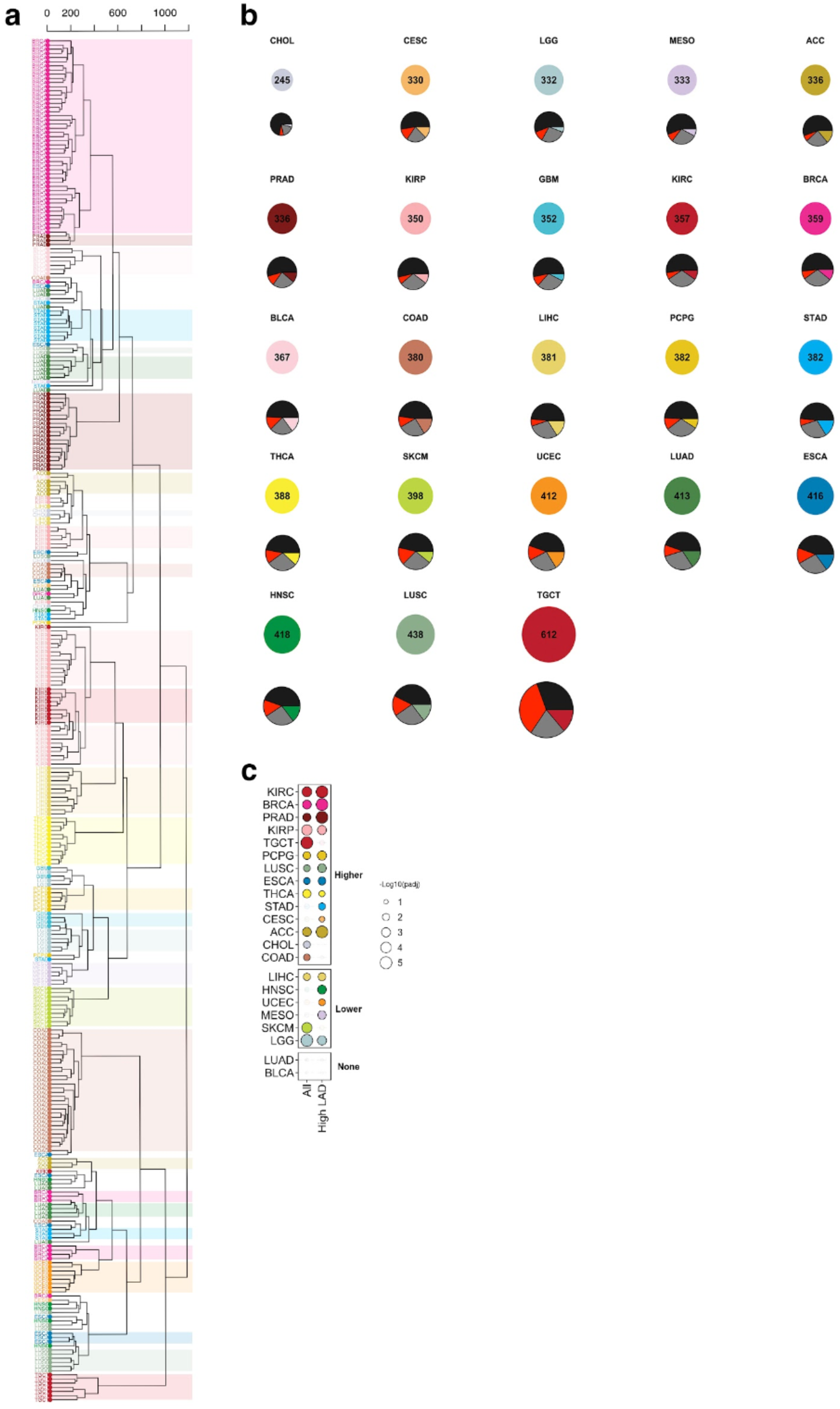
Expansion of RNA polymerase III target genes in human primary tumors. **(a)** Hierarchical clustering of 328 ATAC-seq datasets spanning 23 primary tumor subtypes on the basis of Pol III gene activity signatures. Individual experiments are categorically colored by tumor. **(b)** Overview of the variable breadth and number of tumor-specific Pol III-transcribed gene features defined across all 23 tumor subtypes. Pie charts denote number of predicted “ON” genes in each tumor, and the fractional composition of high uniformity (black), intermediate uniformity (grey), low uniformity (tumor-specific color), and tumor-emergent genes (red). **(g)** Neighborhood analysis of local expression patterns in tumors compared against randomized samples; tumors with significantly higher or lower local expression within a window size of +/-1.5 Mb are colored; permutation p-value.

